# Gonococcal polarization dynamics during adaptation to low oxygen levels

**DOI:** 10.1101/2025.02.18.638858

**Authors:** Paul Schiefer, Berenike Maier

## Abstract

*Neisseria gonorrhoeae* can switch between aerobic and microaerophilic lifestyles, but little is known about the dynamics of this transition. In particular, switching from respiration to partial denitrification is likely to affect the electrophysiology of these bacteria. Here, we use a Nernstian dye to measure the membrane potential of single planktonic cells while they continuously consume the oxygen in the growth medium. We show that cells undergo a characteristic pattern of transient depolarization followed by transient hyperpolarization at a critical oxygen concentration, reminiscent of the response to sudden changes in membrane voltage driven by gated ion-channels. Subsequently, the cells depolarize strongly to a near-constant membrane potential. In the presence of nitrite, the cells repolarize after a delay and the repolarization depends on the oxygen-dependent regulator *fnr* required for denitrification. The temporal behaviour of planktonic cells explains the complex spatio-temporal polarization pattern of the colonies formed by *N. gonorrhoeae*. This sexually transmitted human pathogen experiences different growth environments during transmission. Our results are an important step towards understanding how electrophysiology adapts to changing environments.

## I. INTRODUCTION

Bacteria actively maintain an electrical potential across their cytoplasmic membrane [1, 2]. This potential supports various functions including ATP synthesis, transmembrane transport, and motility [3–5]. Generation of a proton gradient by respiration is crucial for polarization of most bacterial species. Here, we investigate the polarization dynamics during the transition from aerobic to anaerobic respiration.

The bacterial membrane potential has various sources. Active transport of ions and heterogenous permeabilities of ions cause ion gradients across the membrane. Charged macromolecules which are impermeable to the membrane and asymmetry of charges bound on either side of the membrane add to the total electrical potential [1]. Aerobic respiration generates a proton gradient across the membrane whereby oxygen serves as final electron acceptor of the respiratory chain. In the absence of oxygen, many bacterial species employ a denitrification pathway which also generates a proton gradient, yet less efficiently [6]. Electrons are transferred to nitrate or nitrite which are reduced to nitrogen by dedicated denitrification enzymes [6]. *N. gonorrhoeae* is an obligate human pathogen that colonizes mainly the reproductive tract. Thus, it is conceivable that oxygen supply is limited in its natural habitat [7]. It encodes a truncated denitrification pathway that allows it to reduce nitrite to nitrous oxide [7–10]. Under oxygen-limiting conditions the transcription regulator *fnr* is activated. Its regulon comprises genes involved in denitrification including the nitrite reductase *aniA* [11]. Within gonococcal biofilms genes required for denitrification are activated [12] and addition of nitrite enhances biofilm growth [13, 14], indicating that denitrification is important for gonococcal biofilms.

Since the membrane potential is crucial for various functions including ATP production, it was initially thought to be homeostatic. Recently, polarization was investigated at the level of individual bacteria and it was shown to be surprisingly dynamic and heterogeneous [1]. Using genetically encoded probes, *Escherichia coli* cells were shown to blink, i.e. they transiently depolarize [15] or hyperpolarize [16]. Next to genetically encoded probes, Nernstian dyes have been employed for characterizing the dynamics of membrane potential. Using both probes, it was found that the polarization dynamics depend on external factors including growth phase [16], mechanical properties of the environment [17], and antibiotic treatment [18, 19]. In contrast to uncorrelated blinking, colonies of *Bacillus subtilis* show collective polarization dynamics [20, 21]. In these colonies, waves of depolarization travel from the centre to the periphery in an oscillatory fashion. The oscillatory dynamics are based on a potassium channel that is gated by the concentration of a growth resource [20].

Recently, we have discovered collective polarization dynamics in colonies formed by *N. gonorrhoeae* [22]. In liquid, gonococci form spherical microcolonies comprising thousands of cells within minutes [23, 24]. This process of microcolony formation is mediated by type 4 pilus (T4P) driven motility [25, 26], aggregation [27–29], and colony fusion [30, 31]. As colonies reach a critical size, they hyperpolarize at the centre [22]. Subsequently, a shell of hyperpolarization travels radially through the colony and becomes stationary close to the periphery of the colony. In the wake of the hyperpolarized shell potassium influx and slight depolarization was observed. We attribute these spatio-temporal dynamics to the formation of a local oxygen gradient within the colony [22]. It is currently unclear whether the collective polarization dynamics are an emergent property of colonies or whether transient hyperpolarization occurs at a specific oxygen concentration also in planktonic cells.

In this study, we investigate the polarization dynamics of planktonic gonococci as they continuously consume the oxygen in their growth medium. We measure single-cell trajectories of the membrane potential and show that gonococci follow a stereotypical polarization pattern, indicating that transient hyperpolarization occurs not only in colonies but also at the single-cell level. All our data are consistent with the conclusion that hyperpolarization occurs at a specific oxygen concentration. We investigate how gonococci respond to oxygen depletion in the presence of nitrite and show that incomplete denitrification allows gonococci to repolarize after transitioning from aerobic to low-oxygen conditions. Given the complex transmission mechanism of *N. gonorrhoeae*, it is important to understand the electrophysiology of this pathogen in changing environments. Our time-resolved characterization of gonococcal adaptation to anoxic conditions is a crucial step in this direction.

## II. RESULTS

### A. While consuming oxygen, planktonic cells show a synchronous polarization dynamics

In our previous report [22], we showed that when the spherical colonies formed by gonococci reach a certain size, the cells in the centre hyperpolarize and subsequently a shell of hyperpolarized cells travels radially outwards (Fig. 1A, B). We provided evidence that local oxygen gradients form and that their profiles govern the collective polarization dynamics. Our data are consistent with a scenario where all cells hyperpolarize at a critical oxygen concentration, which is present at a specific distance from the edge of the colony (Fig. 1C). Here, we aim to find out whether collective polarization dynamics is an emergent phenomenon of colony formation or whether transient hyperpolarization at a specific (critical) oxygen concentration also occurs in individual cells. To test the second possibility, we characterized the polarization dynamics of non-aggregating gonococci. We chose to work with a *pilE* deletion strain that is unable to form T4P and is therefore immotile and severely impaired in aggregation. We varied the rate of growth resource depletion (including oxygen) by placing the bacteria at varying concentrations into a sealed chamber. We have shown previously, that *N. gonorrhoeae* consume oxygen with a rate that increases with bacterial density in this setup [32]. The membrane potential was determined by quantifying fluorescence of the Nernstian dye TMRM as described in the Methods. If the second hypothesis is correct, then we expect all cells to hyperpolarize simultaneously when the bacteria have depleted the oxygen to the critical concentration (Fig. 1D).

**Fig. 1.**
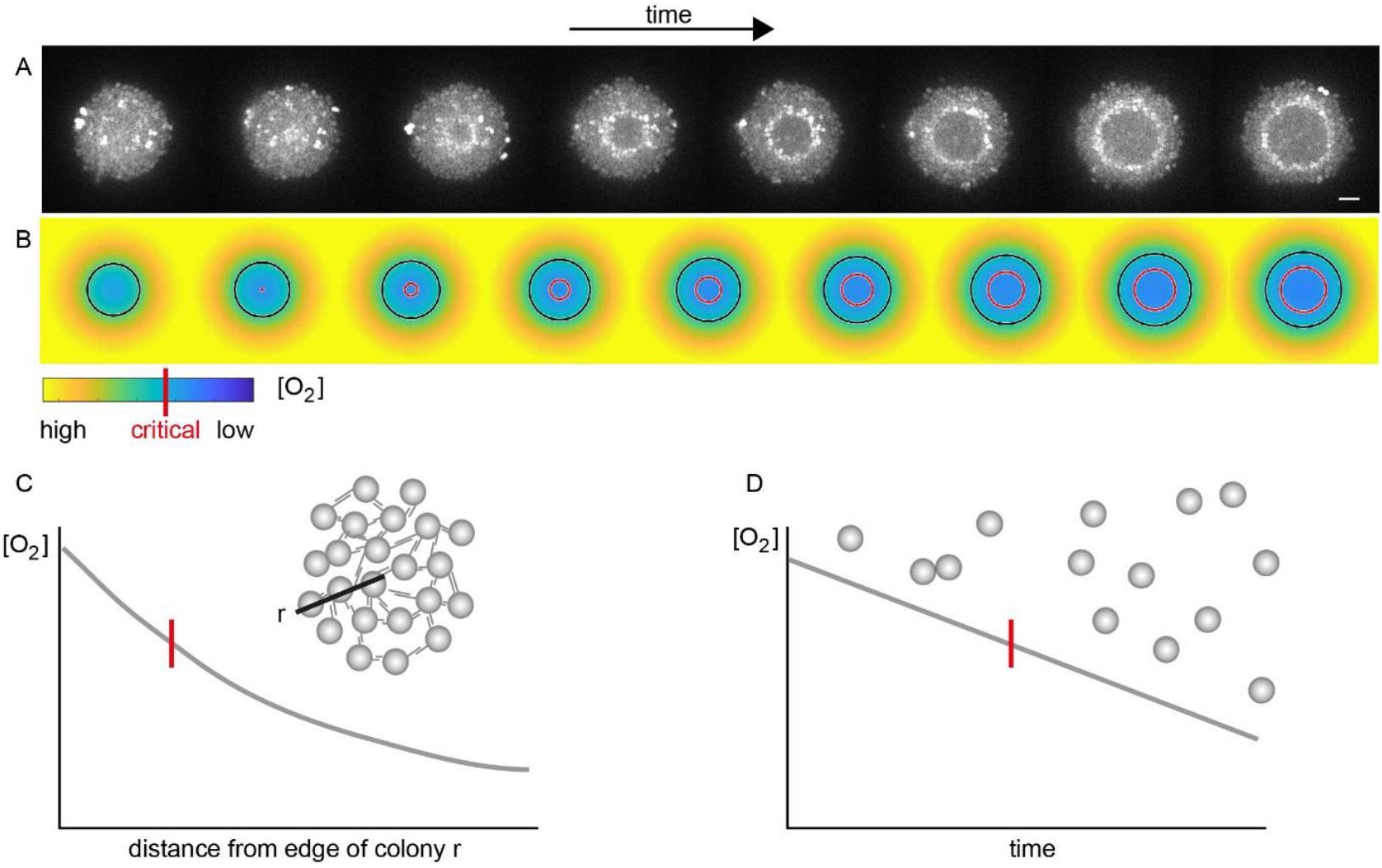
Spatio-temporal and temporal oxygen depletion dynamics. A) Typical dynamics of membrane potential (TMRM fluorescence) of a gonococcal colony. Scale bar: 5 μm. B) Simulation of oxygen concentration. Black circle: edge of colony, red circle: location of critical oxygen concentration. C) Schematic oxygen concentration within a gonococcal colony under continuous oxygen supply. D) Schematic oxygen concentration of a population of planktonic cells in an environment without oxygen exchange. Subpanels A, B were adjusted from [22].

At an inoculation of OD_600_ = 0.03, the initial membrane potential was ≈ −75 *m V* (Fig. 2). After roughly twenty minutes, most cells hyperpolarized simultaneously and transiently to *V*_*m*_ ≈ −90 *mV*. 20-30 min after transient hyperpolarization, the cells depolarized to a value of to a potential of ≈ −40 *mV* (Fig. 2C, Fig. S6). Subsequently, we observed a slow and continuous repolarization to a value of ≈ −50 *mV* (Fig. S6) at 140 min. We note that this phenomenon was observed for each sample, but the exact time at which cells hyperpolarized varied somewhat between samples, most likely due to the fact that the delay between sealing of the sample and start of image acquisition was not measured. Therefore, we show a representative sample in the figures and provide the complete data sets in the Supplementary Information.

**Fig. 2.**
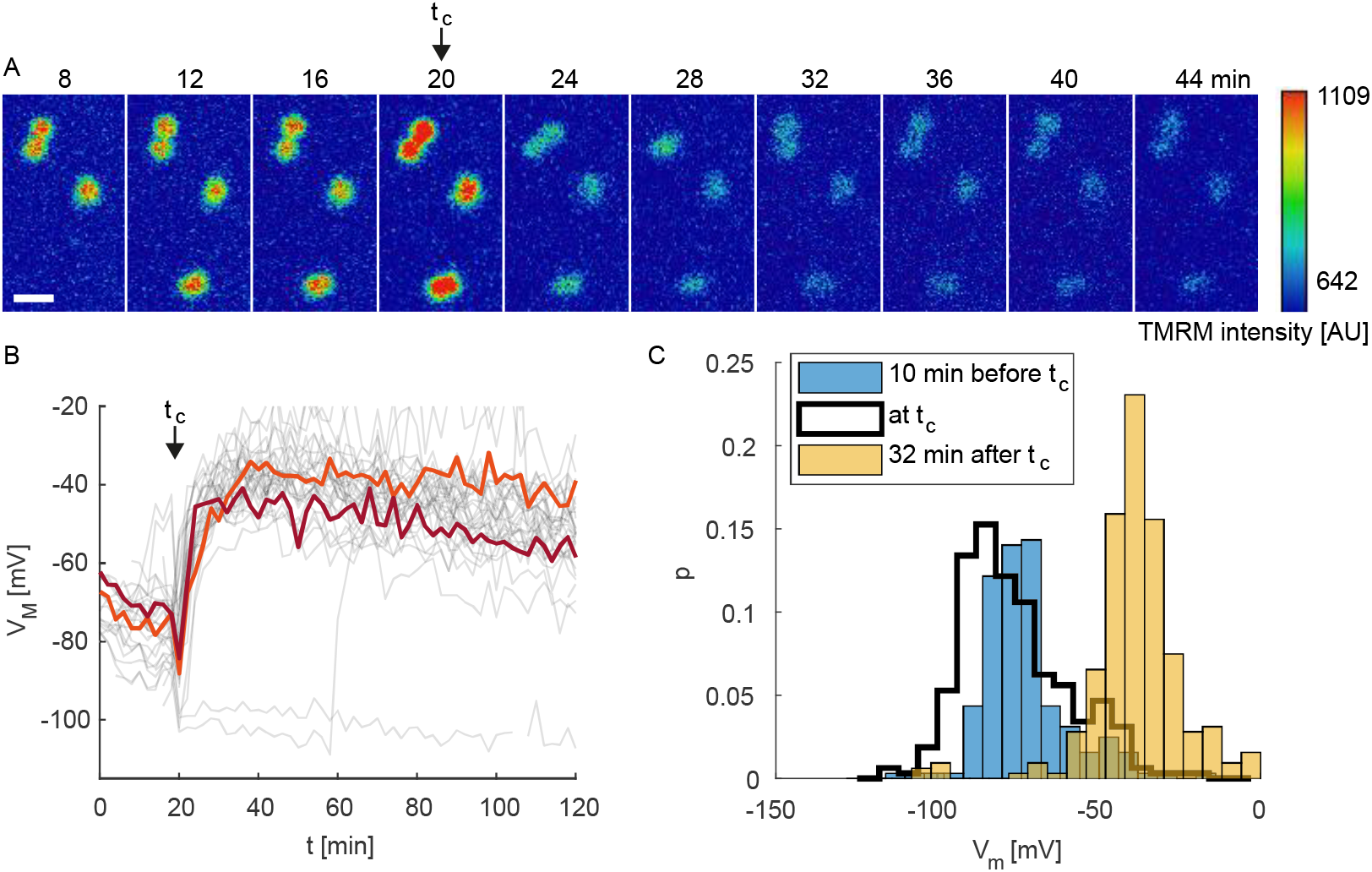
Membrane potential dynamics of individual cells at low level of gas exchange. A) Time lapse of typical TMRM fluorescence at OD_600_ = 0.03. Scale bar: 2 μm. B) Dynamics of membrane potential *V*_*m*_. Two typical trajectories are highlighted for clarity. C) Probability distribution of single cell membrane potential *V*_*m*_ 10 min prior to hyperpolarization (blue bars), during hyperpolarization (black lines), 32 min after hyperpolarization (orange). Data shown in B) result from a single experiment and replicates are shown in Fig. S2C, D. Data shown in C) results from 3 biological culture replicates with N = 109-119 cells per condition.

In most replicates, we found a subset of cells that did not conform to the typical behaviour. A few cells showed very low polarization at the beginning of the experiment. To find out whether theses cells were dead, we co-applied TMRM and the dead stain SytoX. We found that the cells with a membrane potential exceeding ≈ −40 *mV* at the beginning of the experiment showed strong SytoX intensity, indicating that these cells were dead (Fig. S3). Furthermore, a few cells showed stepwise increase SytoX intensity, suggesting that they died during the course of the experiment. Importantly, the fraction of dead cells was below 15 % during the entire experiment. Another small subpopulation of cells showed delayed depolarization after transient hyperpolarization (Fig. 2B, C). We will address this subpopulation later and show that they maintain hyperpolarization via their denitrification pathway.

Next, we addressed the question whether the timing of hyperpolarization was dependent on the rate of growth resource depletion. Therefore, we repeated the experiment at different cell densities. We found that the single cell polarization dynamics were qualitatively comparable to the dynamics found for OD_600_ = 0.03 (Fig. 3A, B). However, decreasing the cell density prolonged the time to transient hyperpolarization and vice versa. The average delay between sealing of the microscopy chamber and transient hyperpolarization, *t*_*c*_, was inversely proportional to cell density (Fig. 3C). This relation indicates that bacteria hyperpolarize at a specific concentration of a growth resource (most likely oxygen [22]).

**Fig. 3.**
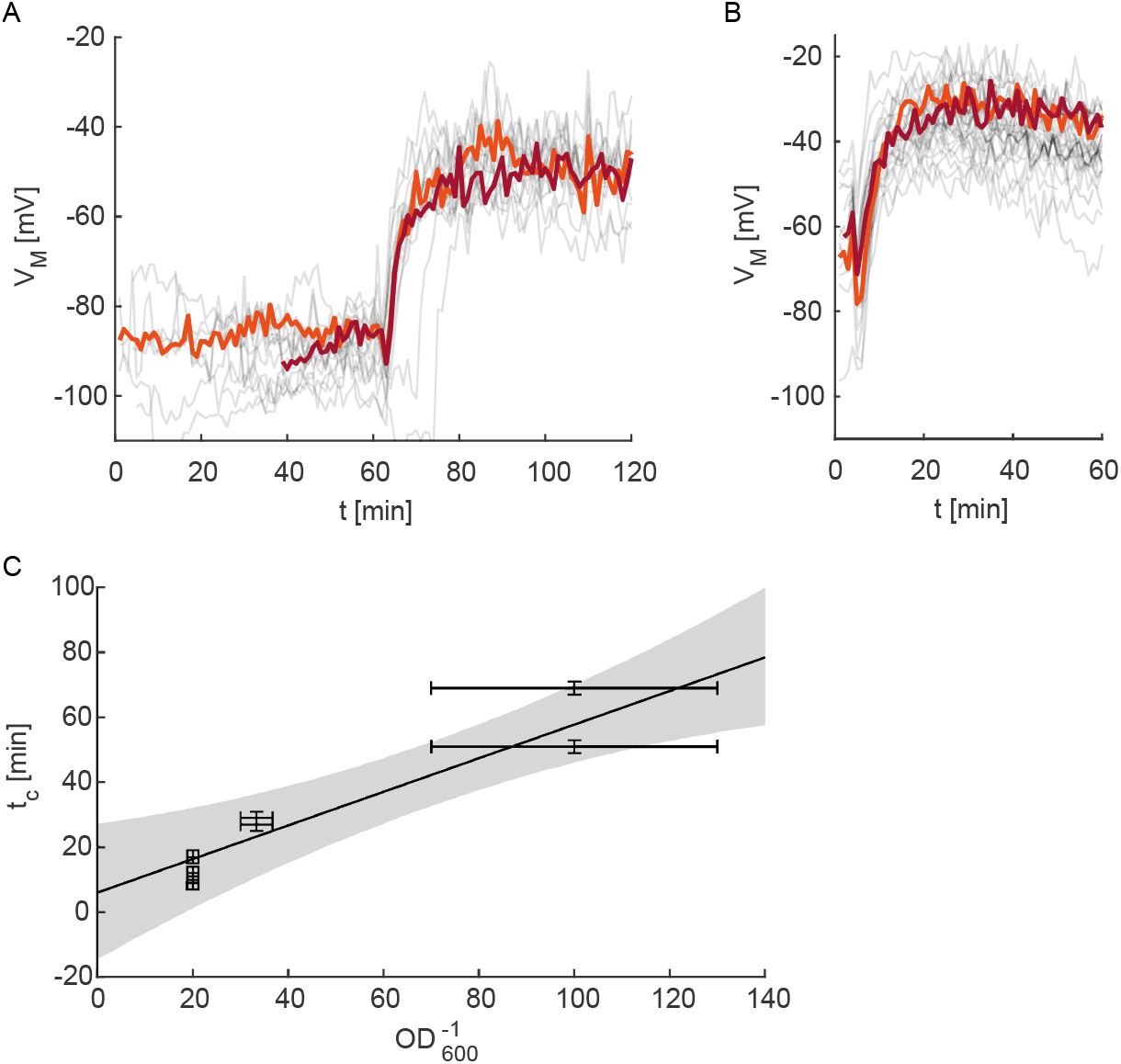
Membrane potential dynamics of individual cells at low level of gas exchange. Dynamics of membrane potential at A) OD_600_ = 0.01, B) OD_600_ = 0.05. C) Time after which the peak occurred, *t*_*c*_, as a function of the inverse OD_600_. Data shown in A, B) result from a single experiment and replicates are shown in Fig. S2. Data shown in C) results from 3 biological culture replicates with N = 48-146 cells per condition.

To ensure that the polarization dynamics were governed by a lack of oxygen and not by a deficiency of another growth resource or chemical communication between cells, we repeated the experiment in a reservoir that allowed the inflow of gas. Under these conditions, no transient hyperpolarization was observed and the membrane potential decreased slowly from an initial value near *V*_*m*_ ≈ −100 *mV* (Fig. 4).

**Fig. 4.**
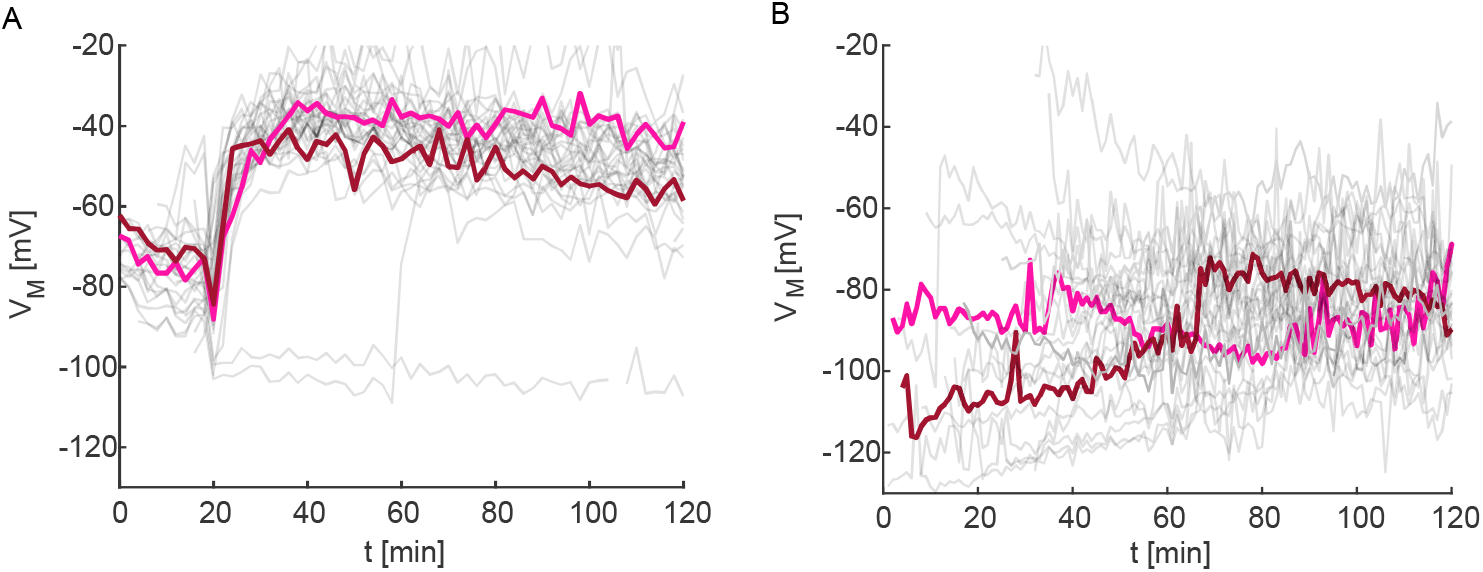
Membrane potential dynamics of individual cells at high level of gas exchange at OD_600_ = 0.03. Trajectories of the membrane potential of single cells in A) air-tight chamber and B) chamber with air-exchange. Data result from a single experiment and replicates are shown in Fig. S4.

We conclude that transient hyperpolarization is not an emergent effect of colony formation, but occurs in response to oxygen depletion in planktonic cells.

### B. Transient depolarization precedes transient hyperpolarization

To characterize the dynamics of transient hyperpolarization in detail, we recorded the trajectories of membrane potential at higher temporal resolution. Single cell trajectories reveal a certain level of heterogeneity (Fig. 5A, Fig. S5-7). In the following, we focus on the dominant behaviour. Most cells transiently depolarized prior to transient hyperpolarization (Fig. 5A, Fig. S5-7). For those cells, the depth of the hyperpolarization event Δ*V*_*M*_ (Fig. 5A) was anticorrelated with the membrane potential before the transient depolarization (Fig. 5B). Furthermore, we found that transient hyperpolarization Δ*V*_*M*_ and width of the transient hyperpolarization peak are independent of the cell concentration (Fig. 5C, D, Fig. S5-7), suggesting that the dynamics of transient hyperpolarization do not depend rate of oxygen consumption. We note that the temporal resolution of membrane potential measurement is limited by the fact that the TMRM dye must equilibrate between the interior and the exterior of the bacterium when the membrane potential changes abruptly. Therefore, we cannot rule out that we are overestimating the widths of the transient depolarization and hyperpolarization events and underestimating the depths of both events. However, if oxygen consumption rate affected the kinetics, we would expect to detect this effect despite the potential broadening of the peaks.

**Fig. 5.**
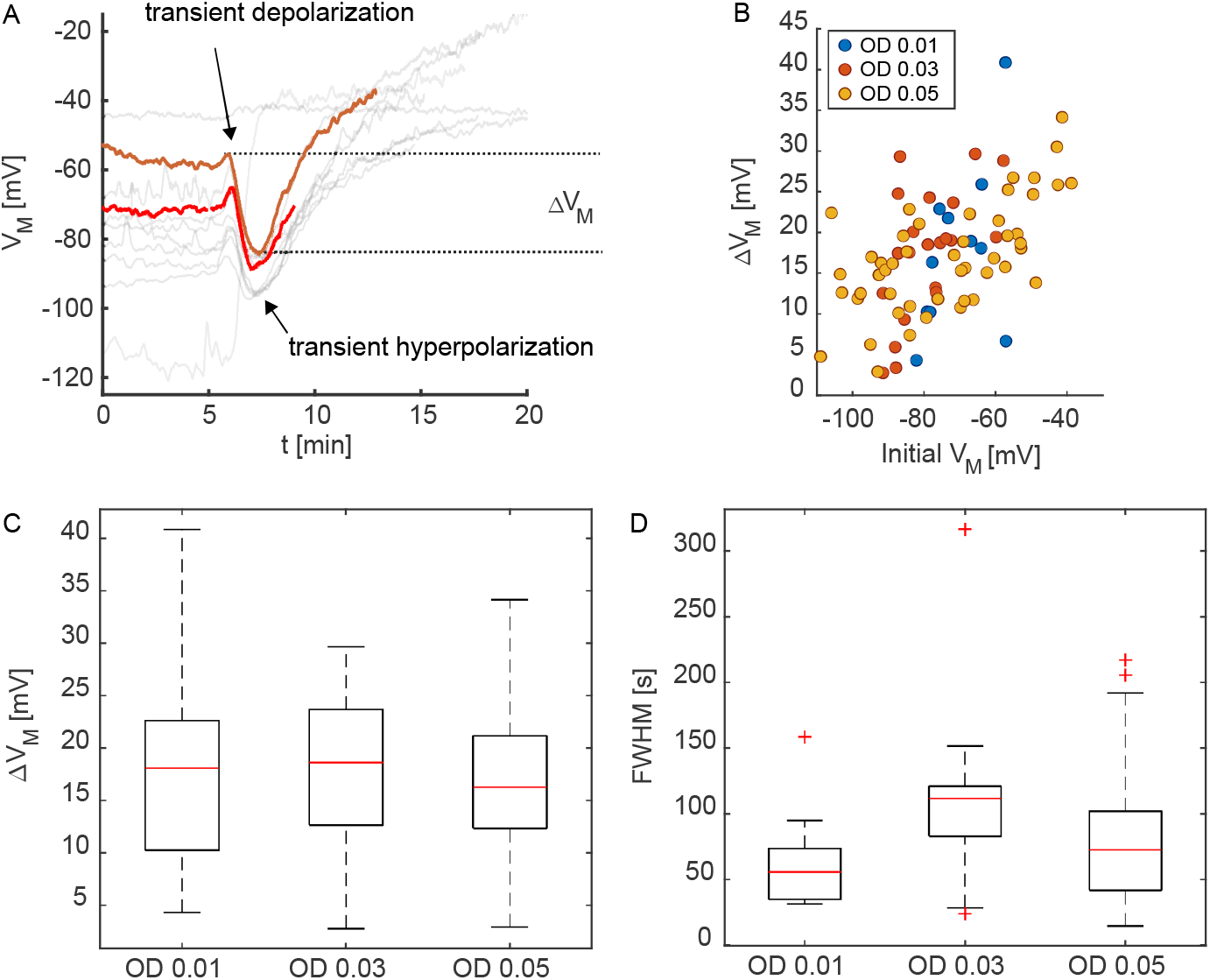
Polarization dynamics close to transient hyperpolarization. A) Single-cell trajectories at OD_600_ = 0.03. B) Correlation between the depth Δ*V*_*M*_ and mean membrane potential prior to transient depolarization. Pearson’s correlation coefficient for combined data *R*_*Pearson*_ = 0.53, *p* = 3.7 · 10^−7^. C) Transient hyperpolarization Δ*V*_*M*_ as a function of OD_600_. D) Peak width (full width at half minimum) as a function of OD_600_. Data shown in A) result from a single experiment and replicates are shown in Fig. S6. Data shown in B-D) results from 3 biological culture replicates with N = 10-48 cells per condition.

To summarize, under conditions of continuous oxygen depletion, the majority of cells show transient depolarization followed by transient hyperpolarization before long-term depolarization.

### C. An alternative electron acceptor of respiratory chain enables recovery of membrane potential

*N. gonorrhoeae* encodes a truncated denitrification pathway that allows it to use nitrite as electron acceptor in the respiratory chain [9, 33] [6, 34]. We assessed how the presence of nitrite affected the polarization dynamics at the single cell level. When we supplemented the medium with 5 mM sodium nitrite, we found that single cells behaved heterogeneously during oxygen depletion. Again, most cells hyperpolarized simultaneously and transiently (Fig. 6A) and subsequently depolarized (Fig. 6A). However, the distribution of the membrane potential of individual cells was shifted to lower values compared to the values in the absence of nitrite supplement (Fig. 6B-E, Fig. S2C-D). Twenty minutes after transient hyperpolarization, the distribution in the presence of nitrite was bimodal because a subpopulation of cells retained the state of hyperpolarization for extended periods of time (Fig. 6D). Thereafter, cells tended to repolarize (Fig. 6C), resulting in a broad distribution of single cell membrane potentials (Fig. 6E).

**Fig. 6.**
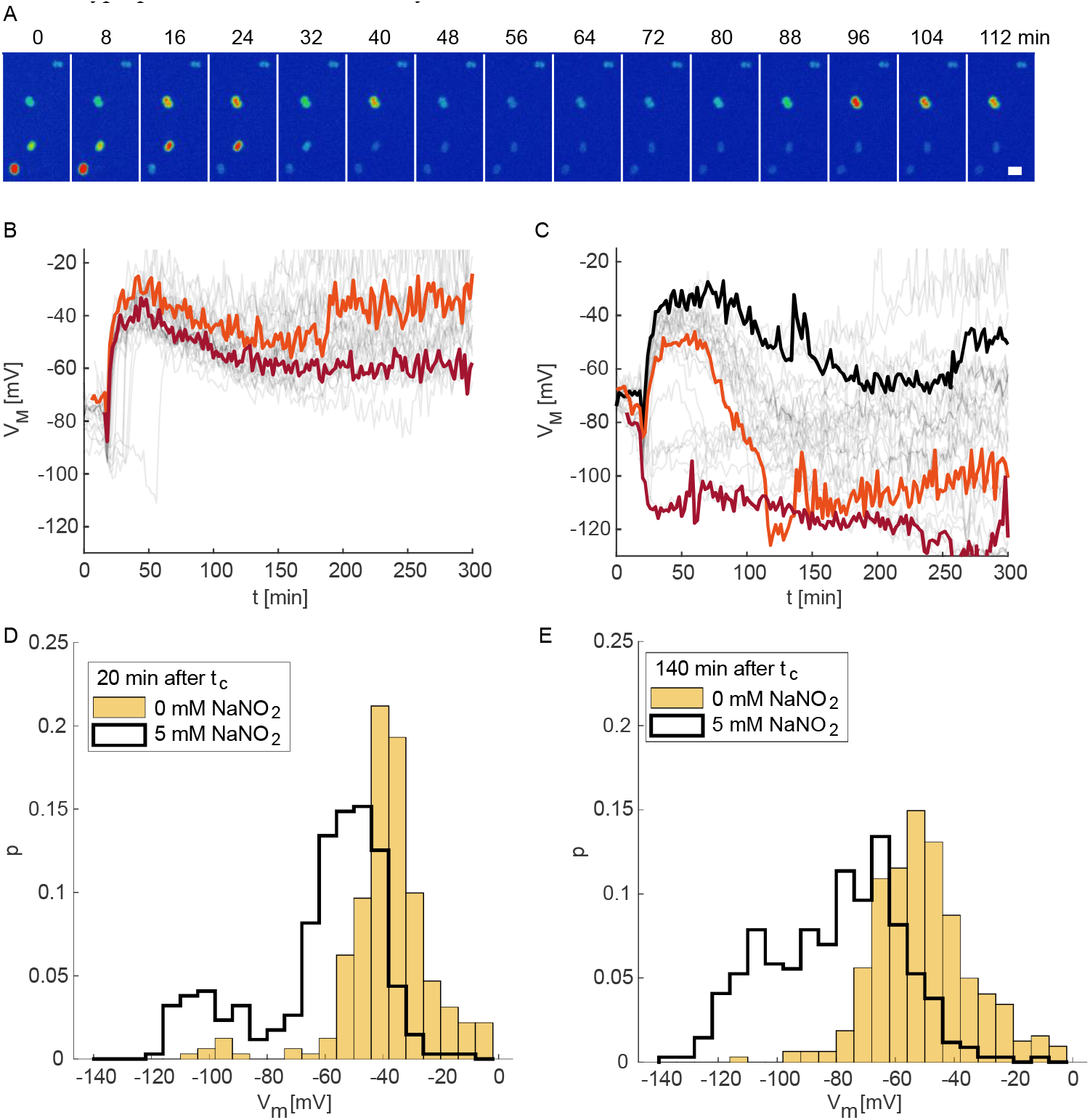
Single cell dynamics of membrane potential with nitrite supplement at OD_600_ = 0.03. A) Time lapse of TMRM fluorescence dynamics. Scale bar: 2 μm. Single cell trajectories of membrane potentials for B) 0 mM NaNO_2_ and C) 5 mM NaNO_2_. Probability distribution of membrane potential without (black line) and with 5 mM NaNO_2_ (orange bars) supplemented D) 20 min after *t*_*c*_ and E) 140 min after *t*_*c*_. Data shown in B, C) result from a single experiment and replicates are shown in Fig. S2C-D and Fig. S8, respectively. Data shown in D, E) results from 3 biological culture replicates with N = 85-126 cells per condition.

The switch from aerobic to anaerobic respiration of *N. gonorrhoeae* is controlled by the fumarate and nitrite reduction regulator *fnr* [11, 34]. Therefore, we speculated that repolarization in the presence of nitrite depends on this regulator. To test this hypothesis, we generated an *fnr* deletion strain and repeated the experiment in the absence (Fig. 7A) and presence (Fig. 7B) of nitrite supplement to the medium. In the absence of nitrite, transient hyperpolarization was visible but less pronounced than in wt* cells (Fig. 7A). After transient hyperpolarization all cells depolarized. In the presence of supplemented nitrite, we observed a similar behaviour (Fig. 7B). Most importantly, we observed no repolarization, demonstrating that upregulation of the denitrification pathway is essential for maintaining prolonged hyperpolarization and repolarization of individual cells.

**Fig. 7.**
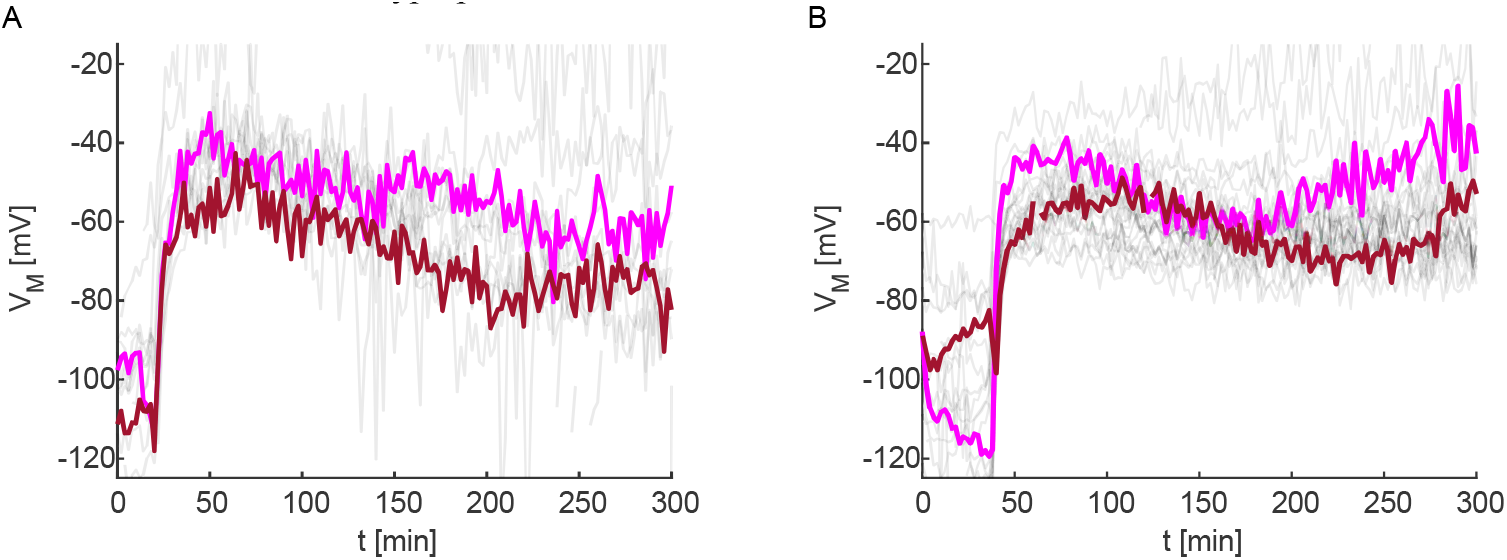
Effect of regulator *fnr* deletion on membrane potential dynamics at OD_600_ = 0.03. Single cell dynamics of membrane potential with medium supplemented with A) 0 mM NaNO_2_ and B) 5 mM NaNO_2_. Data shown result from a single experiment and replicates are shown in Fig. S9.

In summary, denitrification causes repolarization of the cells. The time delay between initial depolarization and repolarization varies between individual cells generating heterogeneity of membrane potentials.

## III. DISCUSSION

In this study, we reveal the characteristic temporal pattern of polarization dynamics during oxygen depletion by *N. gonorrhoeae*. At a critical oxygen concentration, cells show a sequence of transient depolarization and hyperpolarization before depolarizing to a constant value. Our results are most consistent with the following scenario (Fig. 8). at high oxygen concentration, bacteria generate a proton gradient through aerobic respiration (Fig. 8a). Electrons are transferred from NADH to the electron transport chain within the inner membrane. Oxygen serves as the final electron acceptor. This process results in continuous consumption of oxygen in the growth medium. At a critical oxygen concentration, electron transfer slows down and consequently fewer protons are pumped out of the cytoplasm, reducing polarization (Fig. 8B). The cell responds to this depolarization by increasing the polarization by an unknown mechanism (Fig. 8C). As the oxygen levels continue to drop, polarization decreases sharply (Fig. 8D). In this depolarized state, most cells are still alive (Fig. S3) and can repolarize by denitrification (Fig. 7).

**Fig. 8.**
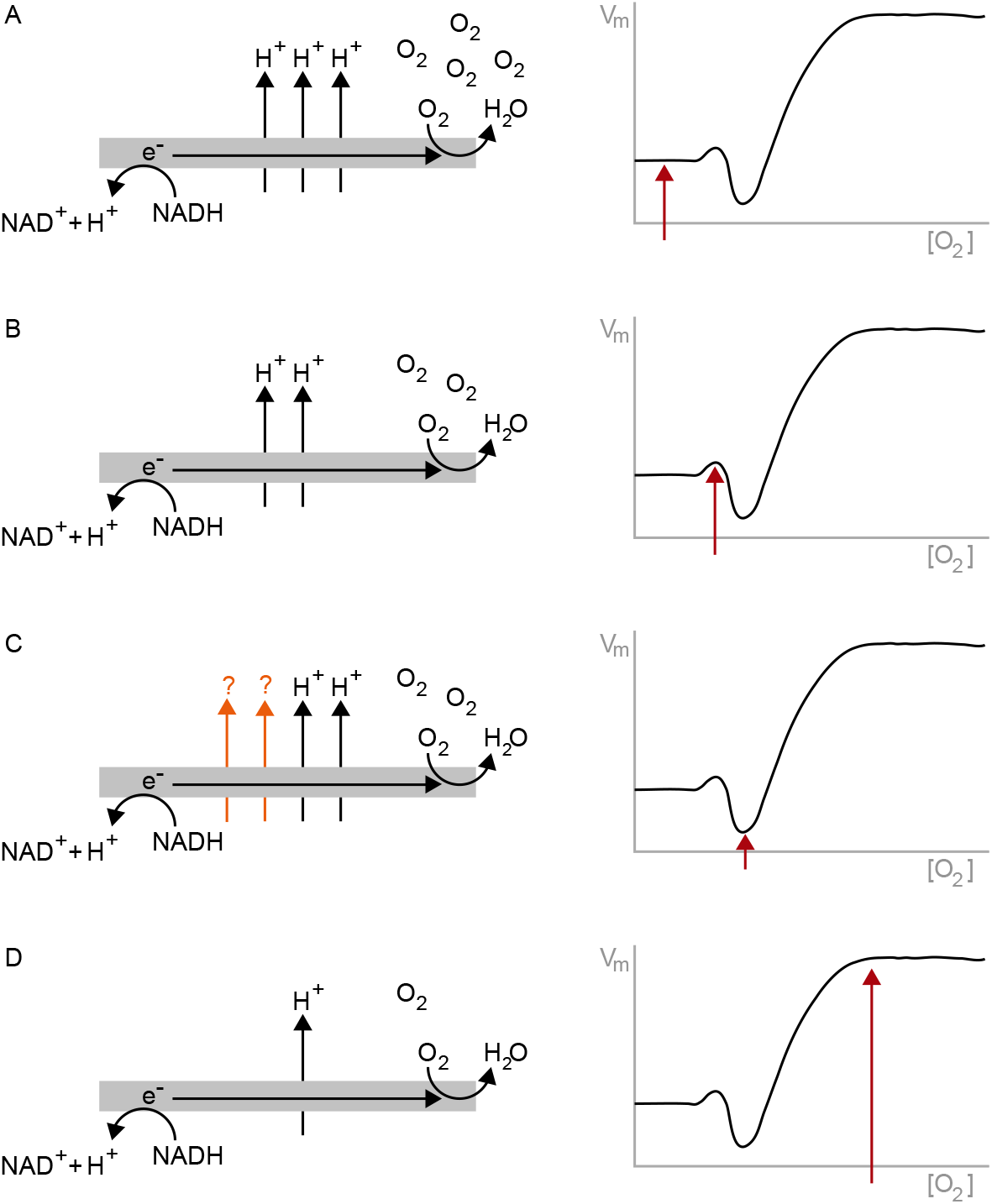
Putative model for polarization dynamics during oxygen depletion. A) At saturating oxygen concentration, cells use the electron transfer chain of aerobic respiration for generating the proton gradient which contributes to the negative membrane potential. B) When the oxygen concentration drops below a critical value, the proton gradient decreases and as a consequence the membrane potential increases. C) Cells respond by transient hyperpolarization through an unknown mechanism. D) Cells depolarize due to reduced respiration.

At the critical oxygen concentration, the characteristic polarization trajectory is reminiscent of the behaviour of *B. subtilis* after transient exposure to high concentration of potassium [20]. In that system, potassium shock resulted in short-term depolarization followed by hyperpolarization. Transient hyperpolarization was caused by potassium efflux through the channel YugO, which is gated by the metabolic state of the cell. In that work, hyperpolarization was considered a cellular response that promotes polarization-dependent uptake of nutrients [20]. In our system, the transient depolarization is most likely caused by reduced proton efflux rather than increased extracellular potassium concentration. However, both affect the membrane potential and lead to the characteristic hyperpolarization response. Mechanistically, a potassium-channel gated by voltage or the metabolic state of the cell could underlie the transient hyperpolarization in the gonococcal system. Although *N. gonorrhoeae* does not possess a *yugO* homologue, it would be interesting to find out which ion channels are involved in the response to oxygen deficiency. Alternatively, transient hyperpolarization could be explained by an increase in NADH production through substrate-level phosphorylation and the TCA cycle [2]. There is also the possibility that the ATP synthase hydrolyses ATP to pump protons out of the cell. This mechanism could explain hyperpolarization during treatment of bacteria with ribosome inhibitors [18]. In contrast to that study, however, we do not expect ATP levels to rise in response to decreased respiratory activity and we consider this explanation to be unlikely.

We estimate the critical oxygen concentration (at which cells depolarize transiently) based on a previous characterization of oxygen consumption by *N. gonorrhoeae* in a slightly poorer growth medium [35]. The saturating oxygen concentration was [*O*_2_]_*inital*_ ≈ 150 ¼*mol l*^−1^ and the oxygen consumption rate was 0.015 *fmol min*^−1^*cell*^−1^. Using this rate, we can estimate the critical oxygen concentration to be [*O*_2_]_*critical*_ ≈ 140 ¼*mol l*^−1^, i.e. roughly 0.9[*O*_2_]_*inital*_. This value is consistent with the critical concentration estimated within gonococcal microcolonies [36].

In this study, we show that *N. gonorrhoeae* depolarize while consuming oxygen from the growth medium. As expected, the cells are able to maintain polarization in the range of ≈ −80 *mV* in the presence of the alternative electron acceptor nitrite. Although the cells were pre-incubated in the presence of nitrite before the transfer to the oxygen-tight chamber, the majority of the population transiently depolarized. After about one hour, most cell repolarized. These dynamics are consistent with various studies showing that genes involved in the truncated denitrification pathway are upregulated under anaerobic conditions [13, 37–39]. We note that the timing of repolarization is heterogeneous and the values of membrane potentials are more broadly distributed than under aerobic conditions.

Radial oxygen gradients form in spherical gonococcal colonies, which change over time [22]. Accordingly, multiple genes involved in oxidative phosphorylation are downregulated while those involved in denitrification tend to be upregulated [38]. In these colonies, transient hyperpolarization waves were observed in the absence, but not in the presence of nitrite [22]. At first glance, this behaviour does not seem to agree with the polarization dynamics of planktonic cells characterized in this study. An important difference in the experimental design of both studies is that the colonies were continuously supplied with fresh medium containing oxygen, and the shape local oxygen gradients changed more slowly. Therefore, cells located at the centre of the colony have enough time to upregulate the denitrification pathway instead of undergoing the period of pronounced depolarization observed here.

## IV. CONCLUSIONS

Gonococcal colonies generate local oxygen gradients that govern the spatio-temporal pattern of polarization dynamics, with a shell of hyperpolarized cells travelling radially through the colony [22]. Here, we show that this spatially localized hyperpolarization observed in colonies can be explained by examining the temporal dynamics of individual planktonic cells during oxygen consumption. We find that transient hyperpolarization occurs immediately following depolarization, suggesting that the cells actively upregulate polarization. In future studies, it will be interesting to characterize the molecular mechanism that causes this polarization overshoot. It is tempting to speculate that transient hyperpolarization serves as signal for transitioning from aerobic to anaerobic metabolism.

## V. METHODS

### A. Culture preparation and microscopy

### 1. Growth media and bacterial strains

Media composition, assay preparation and setup follow [22]. Gonococcal base agar was made from 10 g/L dehydrated agar (BD Biosciences, Bedford, MA), 5 g/L NaCl (Roth, Darmstadt, Germany), 4 g/L K2HPO4 (Roth), 1 g/L KH2PO4 (Roth), 15 g/L Proteose Peptone No. 3 (BD Biosciences), 0.5 g/L soluble starch (Sigma-Aldrich, St. Louis, MO), and supplemented with 1% IsoVitaleX (IVX): 1 g/L D-glucose (Roth), 0.1 g/L L-glutamine (Roth), 0.289 g/L L-cysteine-HCL × H2O (Roth), 1 mg/L thiamine pyrophosphate (Sigma-Aldrich), 0.2mg/LFe(NO3)3 (Sigma-Aldrich), 0.03 mg/L thiamine HCl (Roth), 0.13 mg/L 4-aminobenzoic acid (Sigma-Aldrich), 2.5 mg/L β-nicotinamide adenine dinucleotide (Roth), and 0.1 mg/L vitamin B12 (Sigma-Aldrich). GC medium is identical to the base agar composition but lacks agar and starch. Employed bacterial strains (S1 Table) are derivatives of strain *N. gonorrhoeae* MS11.

### 2. Culture preparation for experiments in sealed microscopy slides

Cells were grown on GC-agar plates overnight at 37°C and 5% CO_2_ for 16 - 20 hours. Bacteria were picked from the plates with inoculation loops and resuspended in GC-medium with 1% IVX, (0.1 - 0.3) μM TMRM, as well as 5 mM NaHCO_3_ for additional CO_2_. We also added 2% MilliQ H_2_O as this step was included in the preparation process described by Hennes *et al*. [22]. In experiments with sodium nitrite we added 5mM of NaNO_2_ to the medium. The cell suspension was vortexed for 2 minutes and adjusted to the desired OD_600_. Cultures were then placed in a shaking incubator for 30 minutes at 37°C and 5% CO_2_. Subsequently, 50 μl of the cell suspension were inoculated between a microscopy glass and a glass coverslip. The edges of the coverslip were sealed with a mixture of equal amounts Vaseline, wool fat and Paraplast (Roth) to prohibit gas exchange.

#### 3. Confocal imaging

Cells were imaged using an inverted microscope (Ti-E Nikon) equipped with a CSU-X1 (Yokogawa) spinning disk unit, a 100x CFI Apo Tirf objective (Nikon), an EMCCD camera (iXon 897 X-11662, Andor) and 488 nm and 561 nm excitation lasers.

### B. Determination of single cell potential

Videos were prepared for further analysis in ImageJ. For each field of view translational motion was corrected via the plug-in HyperStackReg. The resulting data was saved as TIFF image series for the respective channels. Cell positions and tracks were determined using the tracking plug-in TrackMate. We wrote MATLAB scripts for obtaining the fluorescence intensity of each tracked cell for each point in time. For further analysis and presentation, we filtered the tracks according to their length. For experiments with low frame rates of 1–2 frames per minute, tracks with a length of less than 60% of the total recording time were discarded; for high-frequency measurements with 2–5 frames per second, tracks with a length of less than 30% of the total recording time were discarded. This filtering was applied to remove erroneously connected points during tracking and tracks from cells that settled late in the experiment when the transient hyperpolarization had already occurred.

We wrote MATLAB scripts to analyze the width and height of the transient hyperpolarization in tracks recorded at 2–5 frames per second, and only considered tracks in which the transient hyperpolarization was present. This was assessed manually and individually for each track. In total, we analyzed 10/19 cell tracks from the OD_600_ = 0.01 experiments, 22/42 tracks from the OD_600_ = 0.03 experiments, and 48/90 tracks from the OD_600_ = 0.05 experiments. To reduce noise, we smoothed the data from high frame rate experiments by applying a 10-second window sliding average to the tracks.

As TMRM is a Nernstian dye, its distribution follows Boltzmann’s law 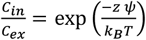 with the ratio of internal to external dye concentration 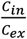, the charge of the dye *z*, the potential difference between the interior and the outside of the cell *ψ*, the Boltzmann constant *k*_*B*_ and the temperature *T*. However, experimentally we can only obtain the ratio of fluorescence 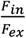 inside and outside the cell rather than the dye concentration itself. Due to the point spread function (PSF) of the microscope setup we underestimate the true ratio of 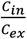 by simply measuring 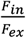. We calculated a correction factor *S(F*_*in*_*/F*_*ex*_) to correct for the effect of the PSF. This function was derived from a three-dimensional convolution model following the procedure described recently [40]. The PSF of our microscope was determined as the pixel intensities of fluorescent beads with a diameter of 100 nm (TetraSpeck™, Thermo Fisher) that are fixed on a glass coverslip. To obtain the three-dimensional PSF, we scanned the fluorescence of the beads in z-direction in a range of ±1.62 μm in 0.079 μm intervals. Three beads were characterized and the resulting intensity values averaged. The background was subtracted and the resulting intensities were normalized. To obtain the correction factor *S(F*_*in*_*/F*_*ex*_ ) (Fig. S1), we simulated an idealized *N. gonorrhoeae* cell on a microscopy glass. We modelled a cubic box with length, width and height of 10 μm. At the centre of the x-y-plane we placed a cell in form of a sphere with radius 0.5 μm for which the bottom part is in contact with the box. The rest of the box is filled with medium. For the dye concentration we used *C*_*i*_*(x*_*i*_, *y*_*i*_, *z*_*i*_) = 0 for *z* ≤ 0, as this corresponds to the glass in our experiment. Inside of the cell, the concentration is set to *C*_*i*_*(x*_*i*_, *y*_*i*_, *z*_*i*_) = *C*_*in*_. In the medium, the concentration of pixels is set to *C*_*i*_*(x*_*i*_, *y*_*i*_, *z*_*i*_) = *C*_*ex*_. From this model, we obtained the simulated image intensities *I*_*m*_*(x*_*j*_, *yj, z*_0_) at the central focal plane of the cell *z*_0_ = 0.5 μm by convolution of the dye concentrations and the PSF. As we know the exact position of the cell, we are able to calculate the internal fluorescence.

## C. Text editing

The software DeepL was used to improve the text flow.

## ACKNOWLEDGMENTS

We thank Marc Hennes for supporting the start of this project and the Maier lab for stimulating discussions. This work was supported by the Deutsche Forschungsgemeinschaft through grant SPP2389 MA3898 and the Center for Molecular Medicine Cologne through grant B06.

## Notes

### Competing Interest Statement

The authors have declared no competing interest.

