## Supplemental Information for "Gonococcal polarization dynamics during adaptation to low oxygen levels"

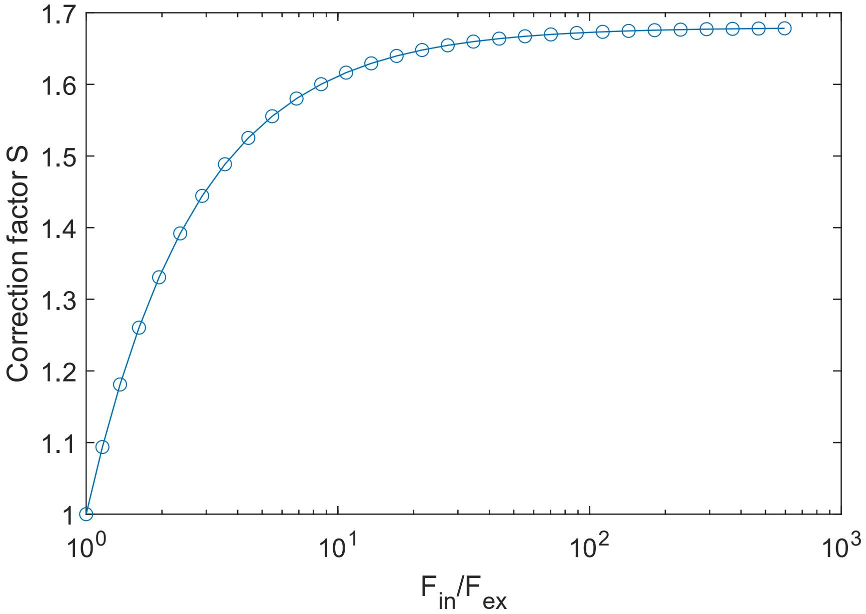


Fig. S1 Correction factor for the ratio between internal and external TMRM concentration.

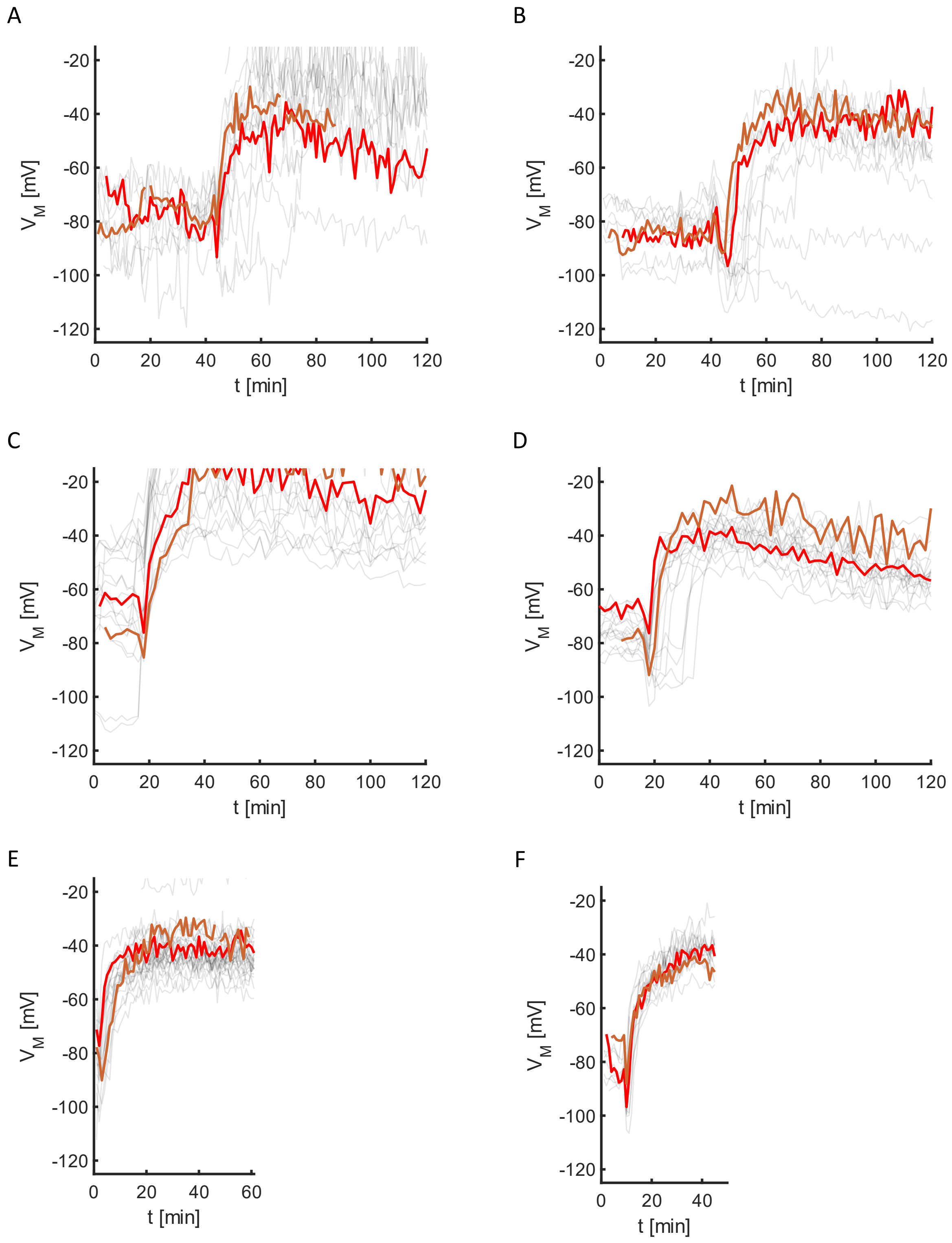


Fig. S2 Membrane potential dynamics of individual cells in air-tight chambers. Biological culture replicates of data shown in A-B) Fig. 3A at $OD_{600}$ = 0.01, C-D) Fig. 2B at $OD_{600}$ = 0.03, E-F) Fig. 3B at $OD_{600}$ = 0.05. Two representative trajectories are highlighted in each graph.

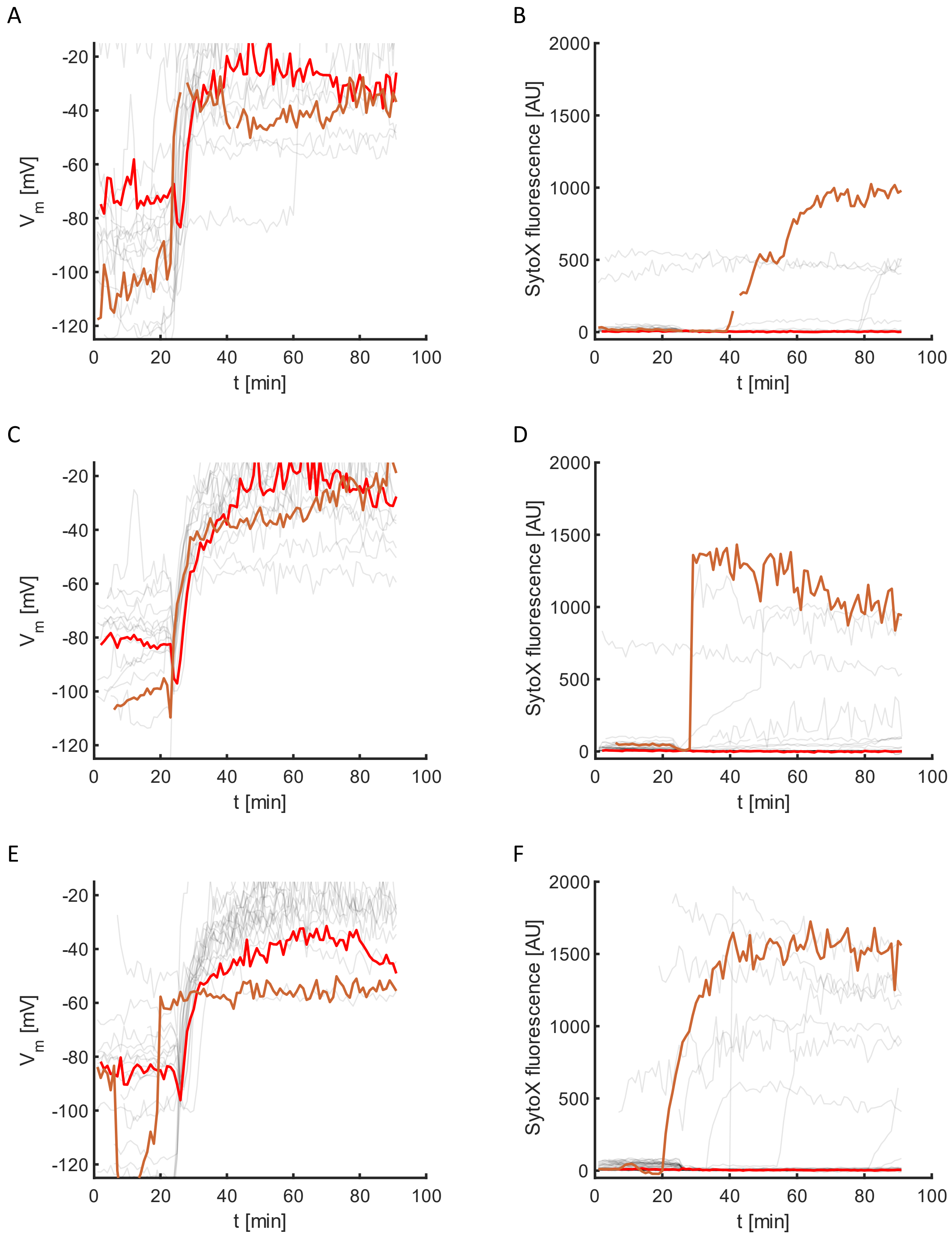


Fig. S3 Correlation between polarization dynamics and cell death in air-tight chambers at $OD_{600}$ = 0.05. A), C), E) Membrane potential of individual cells. B), D), F) Fluorescence signal of the dead stain SytoX. Neighbouring graphs belong to the same day of experiment (A-B), C-D), E-F)). Two cells are highlighted for each experiment; one that dies over the course of the experiment and one that stays alive. While almost all cells strongly depolarize after about 20 minutes only a small fraction of cells dies. At $t$ = 5 min a total of 3 out of 111 tracked cells are dead, at the time of the hyperpolarization 7 out of 145 tracked cells are dead and at $t$ = 90 min 17 out of 126 tracked cells are dead.

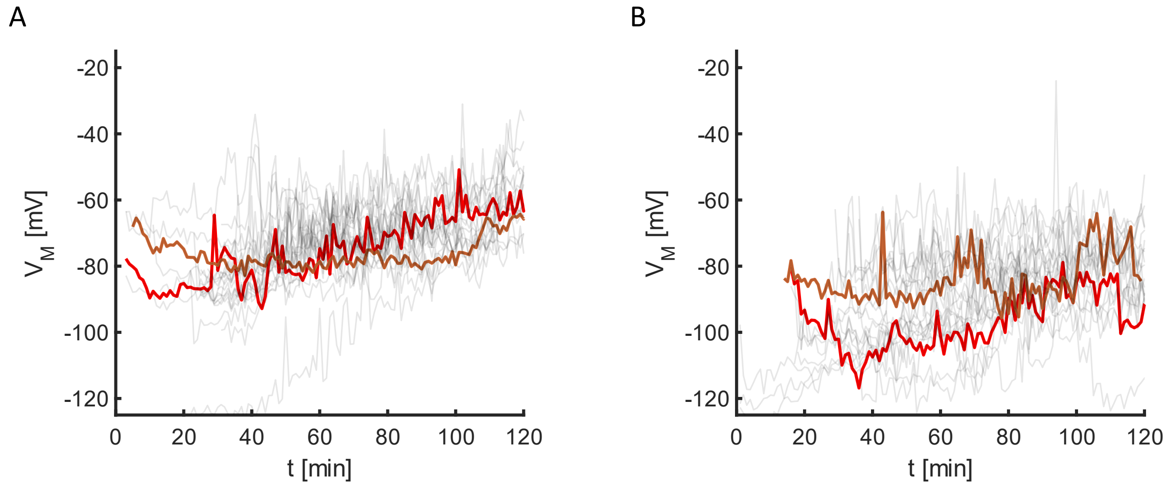


Fig. S4 Membrane potential dynamics of individual cells in chambers with air-exchange at $OD_{600}$ = 0.03. A, B) Biological culture replicates of data shown in Fig. 4B. Two representative trajectories are highlighted in each graph.

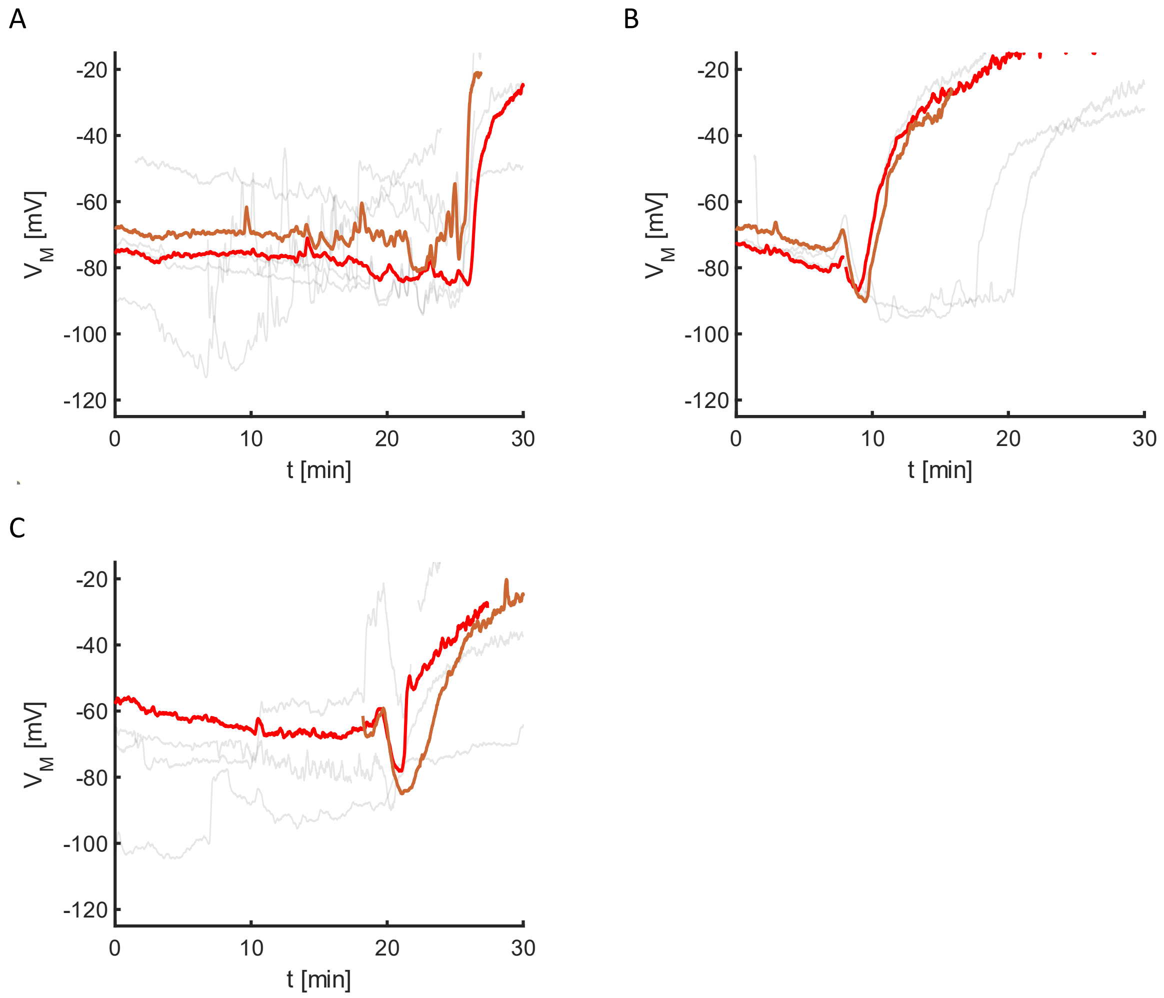


Fig. S5 Polarization dynamics close to transient hyperpolarization in air-tight chambers at $OD_{600} = 0.01$. A-C) show three biological culture replicates. Two representative trajectories are highlighted in each graph. Image acquisition started 22 minutes after sealing the microscopy chambers.

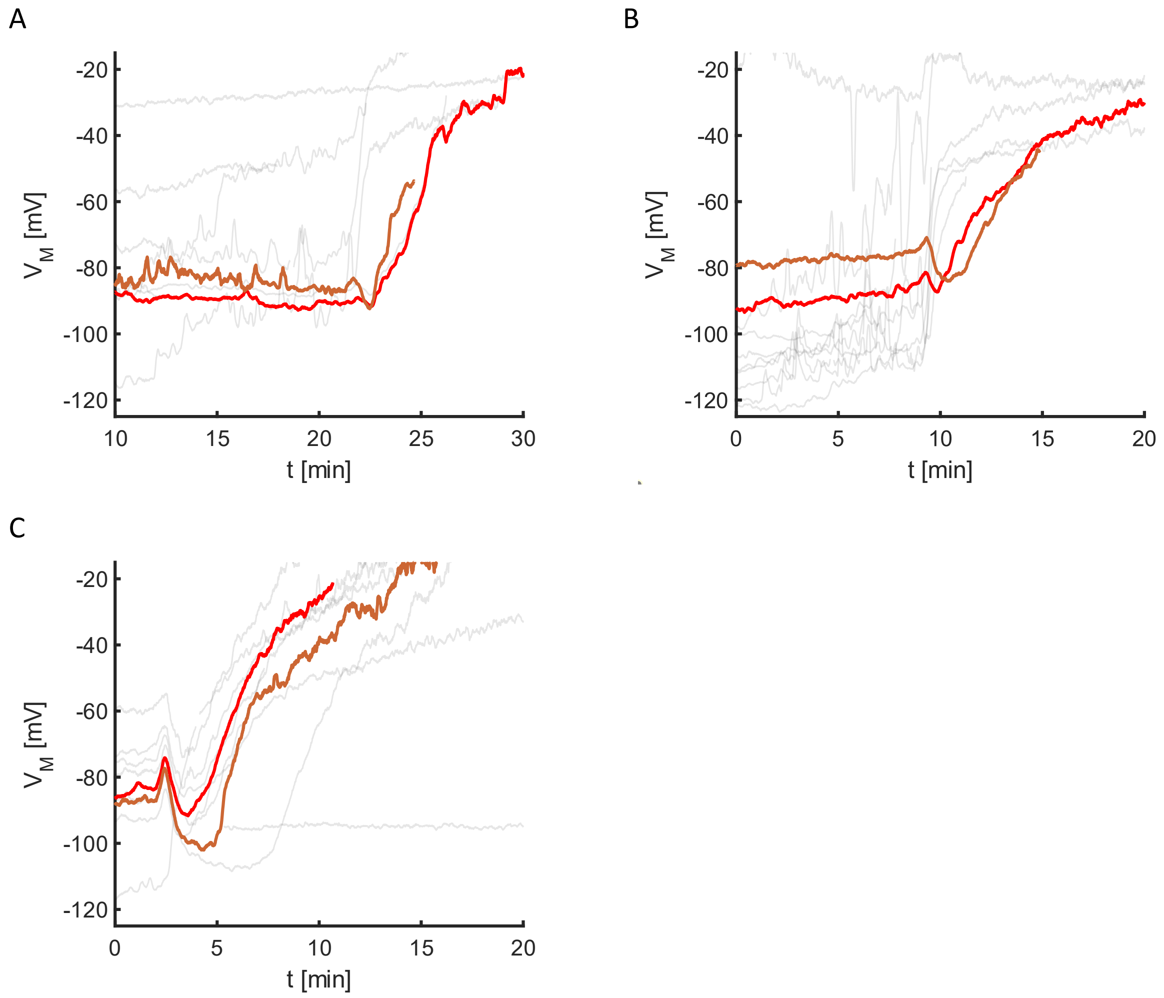


Fig. S6 Polarization dynamics close to transient hyperpolarization in air-tight chambers at $OD_{600}$ = 0.03. A-C) show three biological culture replicates. Two representative trajectories are highlighted in each graph. Image acquisition started 12 minutes after sealing the microscopy chambers.

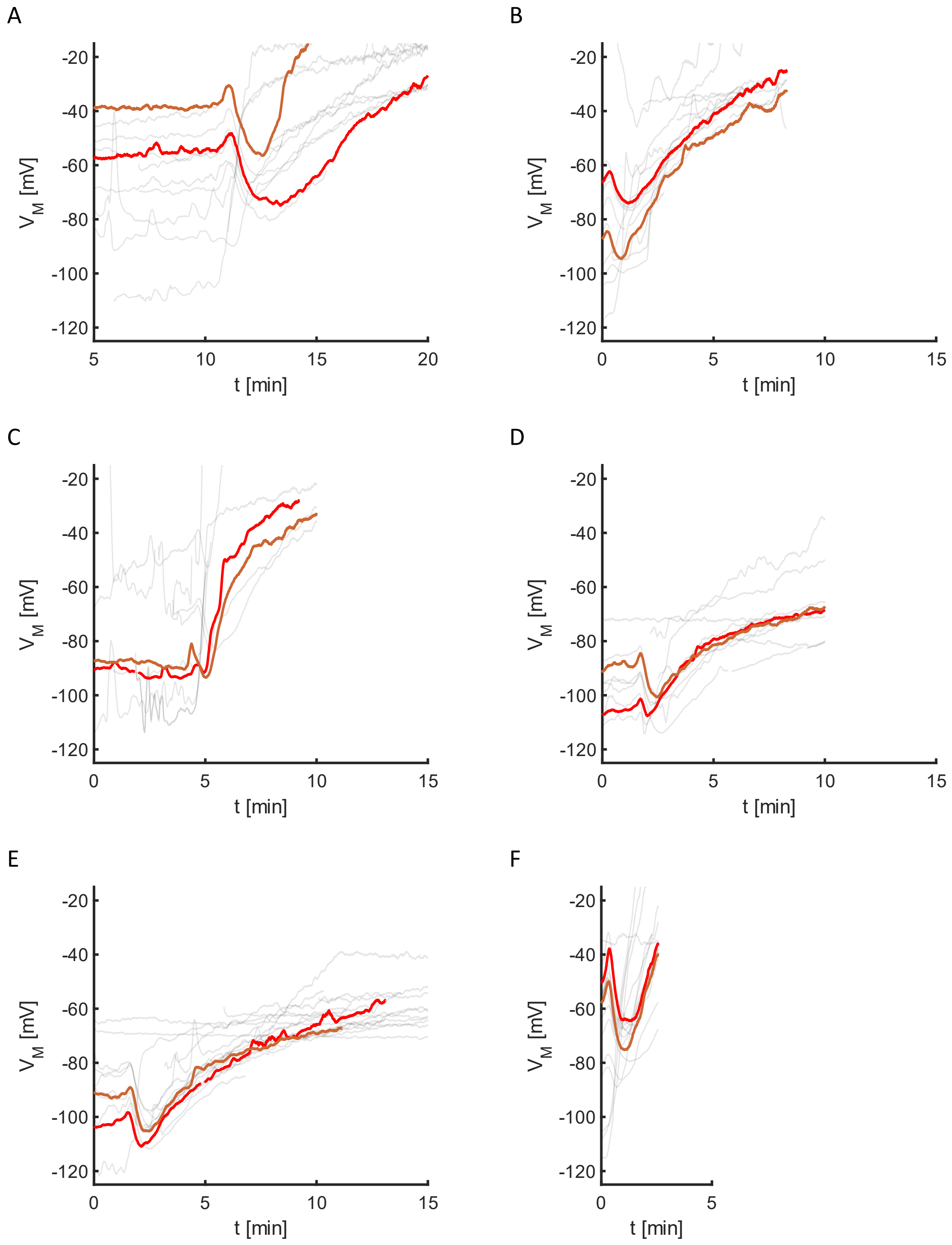


Fig. S7 Polarization dynamics close to transient hyperpolarization in air-tight chambers at $OD_{600}$ = 0.05. A-F) show four biological culture replicates and two technical replicates. Two representative trajectories are highlighted in each graph. Image acquisition started 2 minutes after sealing the microscopy chambers.

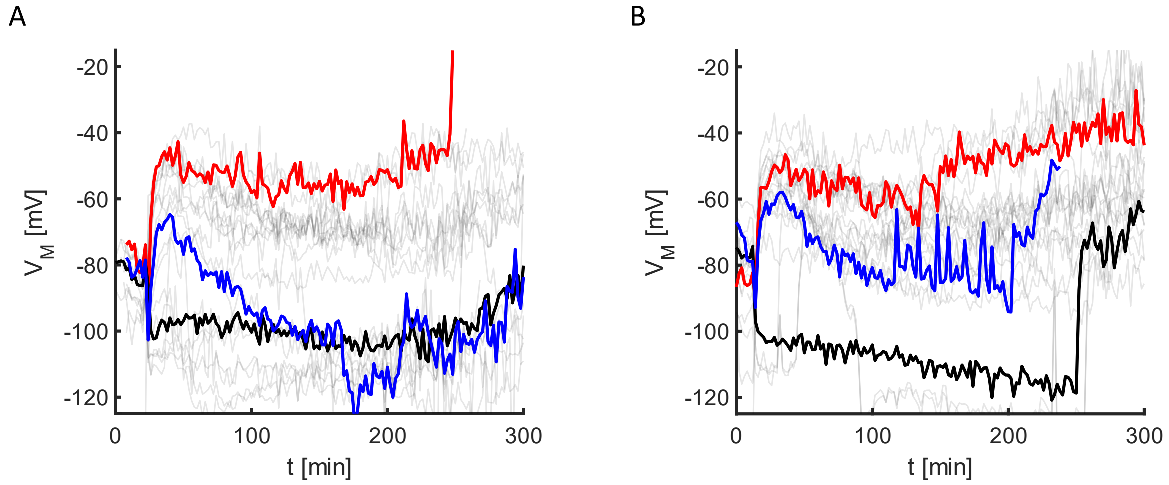


Fig. S8 Single cell dynamics of membrane potential with nitrite supplement (5 mM $NaNO_2$) in air-tight chambers at $OD_{600}$ = 0.03. A-B) Biological culture replicates of data shown in Fig. 6C. Three typical trajectories are highlighted.

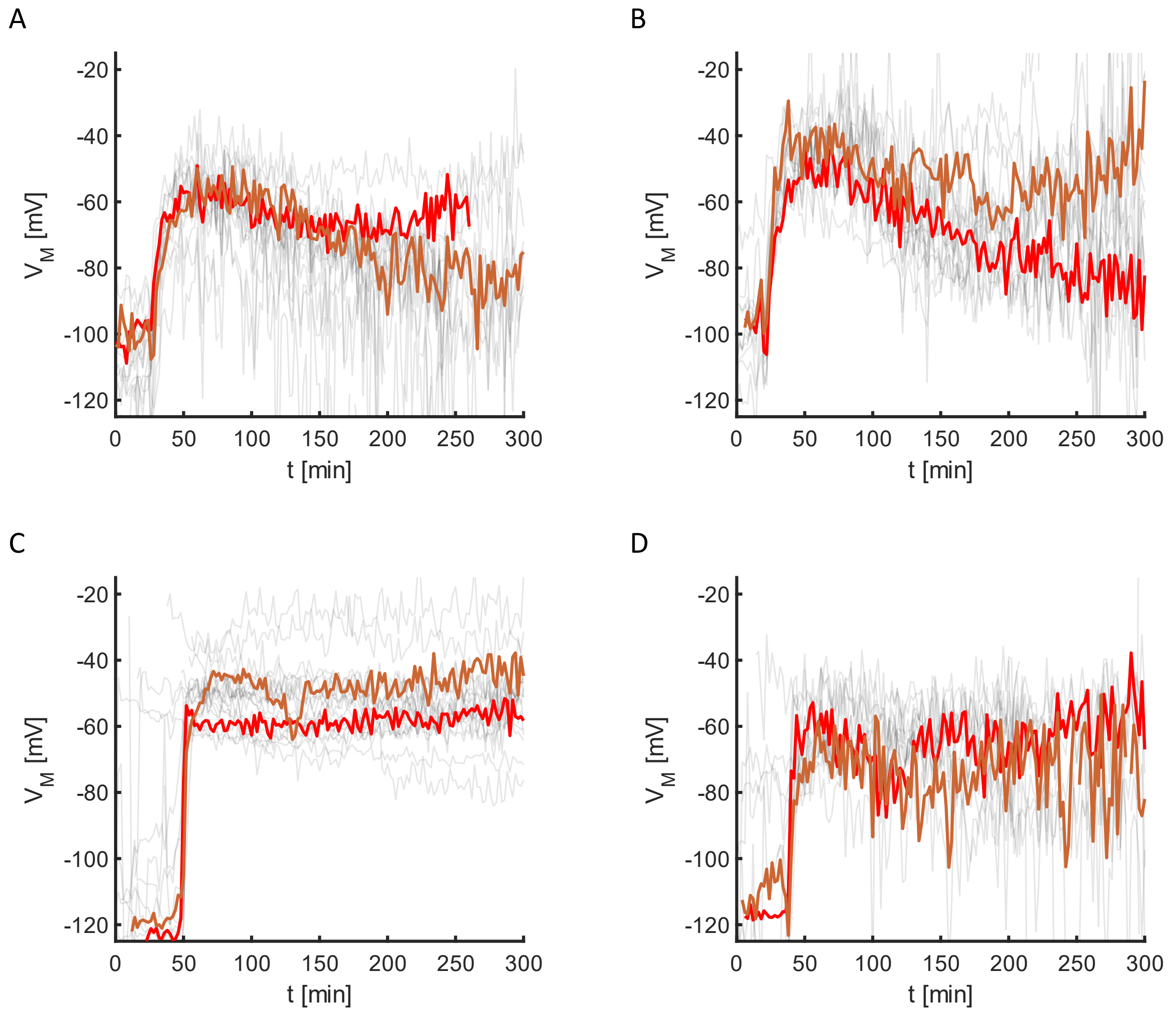


Fig. S9 Effect of regulator *fnr* deletion on membrane potential dynamics in air-tight chambers at $OD_{600}$ = 0.03. A, B) Biological culture replicates of data shown in Fig. 7A at 0 mM $NaNO_2$. C, D) Biological culture replicates of data shown in Fig. 7B at 5 mM $NaNO_2$. Two typical trajectories are highlighted.

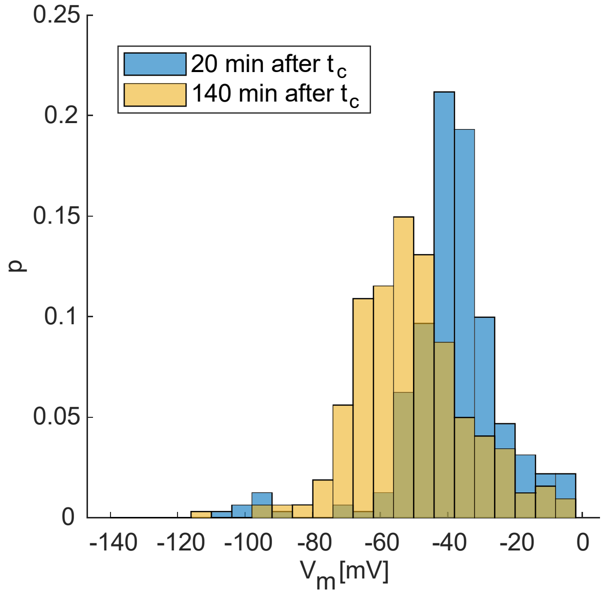


Fig. S10 Probability distribution of single cell membrane potential at $OD_{600}$ = 0.03. 20 min (orange bars) and 140 min (blue bars) after hyperpolarization.

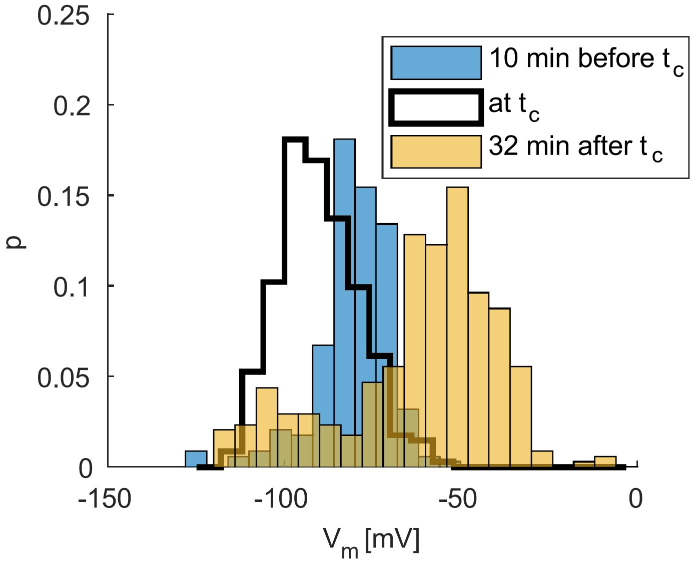


Fig. S11 Probability distribution of single cell membrane potential with 5 mM $NaNO_2$ supplement at $OD_{600}$ = 0.03. 10 min prior to hyperpolarization (blue bars), during hyperpolarization (black line), 32 min after hyperpolarization (orange bars) with 5 mM $NaNO_2$ supplemented.

| Strain | Relevant genotype | Source/Reference |
|---|---|---|
| *wt** (Ng150) | *G4::aac* | [1] |
| *ΔpilE* (Ng196) | *pilE::cat*<br>*G4::aac* | This study, [2] |
| *ΔpilE Δfnr* (Ng287) | *fnr::kan*<br>*pilE::cat*<br>*G4::aac* | This study, [2] |

Table S1 Bacterial strains used in this study.

1. Zollner, R., Cronenberg, T., Kouzel, N., Welker, A., Koomey, M., and Maier, B. (2019). Type IV Pilin Post-Translational Modifications Modulate Material Properties of Bacterial Colonies. Biophys J *116*, 938-947. 10.1016/j.bpj.2019.01.020.

2. Welker, A., Cronenberg, T., Zollner, R., Meel, C., Siewering, K., Bender, N., Hennes, M., Oldewurtel, E.R., and Maier, B. (2018). Molecular Motors Govern Liquidlike Ordering and Fusion Dynamics of Bacterial Colonies. Phys Rev Lett *121*, 118102. 10.1103/PhysRevLett.121.118102.
